# A Spatial Transcriptomics Study of the Brain-Electrode Interface in Rat Motor Cortex

**DOI:** 10.1101/2021.12.03.471147

**Authors:** Quentin A. Whitsitt, Bella Patel, Brad Hunt, Erin K. Purcell

**Affiliations:** Department of Biomedical Engineering, Michigan State University. East Lansing, Michigan, USA; Department of Electrical and Computer Engineering, Michigan State University. East Lansing, Michigan, USA

## Abstract

The study of the foreign body reaction to implanted electrodes in the brain is an important area of research for the future development of neuroprostheses and experimental electrophysiology. After electrode implantation in the brain, microglial activation, reactive astrogliosis, and neuronal cell death create an environment immediately surrounding the electrode that is significantly altered from its homeostatic state. To uncover physiological changes potentially affecting device function and longevity, spatial transcriptomics was implemented in this preliminary study to identify changes in gene expression driven by electrode implantation. This RNA-sequencing technique (10x Genomics, Visium) uses spatially coded, RNA-binding oligonucleotides on a microscope slide to spatially identify each sequencing read. For these experiments, sections of rat motor cortex implanted with Michigan-style silicon electrodes were mounted on the Visium slide for processing. Each tissue section was labeled for neurons and astrocytes using immunohistochemistry to provide a spatial reference for mapping each sequencing read relative to the device tract. Results from rat motor cortex at 24 hours, 1 week, and 6 weeks post implantation showed up to 5811 differentially expressed genes between implanted and non-implanted tissue sections. Many of these genes are related to biological mechanisms previously reported in studies of the foreign body response to implanted electrodes, while others are novel to this study. These results will provide a foundation for future work to both improve and measure the effects of gene expression on the long-term stability of recordings from implanted electrodes in the brain. Ongoing work will expand on these initial observations as we gain a better understanding of the dynamic, molecular changes taking place in the brain in response to electrode implantation.

## I. Introduction

Intracortical microelectrode implants have the capacity to stimulate nervous tissue and/or record electrical signals from the brain. This functionality allows for a wide range of applications, from treating debilitating neurological diseases to preclinical and basic neuroscientific research. Clinical applications of these devices have progressed rapidly over the past decade and have recently gained newfound interest due to the potential for use in brain-computer interfaces (BCIs). Recent advances in recording-based implants have restored quadriplegic^1,2^ or quadriparesis^3^ patients’ ability to communicate. BCI research has shown the capability of recording implants in motor cortex to drive movement of a robotic limb or computer cursor^4,5^. Generally, intracortical electrodes are used in these applications to record electrical signals from specific areas of the brain. An algorithm is then trained to decode the electrical signals into meaningful information that can be used to drive the BCI and produce the desired outcome.

Limitations to the long-term use of recording implants *in vivo* include observations of a decrease in signal quality over time^6–10^ as well as signal instability^6,10–12^. Decline of signal quality is characterized by decreased amplitude of unit waveforms, number of resolvable units, signal-to-noise ratio, and an increase in impedance. All of these trends can contribute to a decline in device function by reducing information transfer at the brain-electrode interface^13^. This creates limitations for current BCIs because loss of information directly affects decoder performance^13^, and it also creates limitations in research settings when trying to study complex and precise neurological activity over a long period of time.

There are multiple possible mechanisms that may lead to signal quality and stability decline^7,14^. In addition to mechanical/electrical failure and micromotion, a long-standing line of inquiry into the origins of signal decay focuses on the biological response to implants in the brain, known as the foreign body reaction (FBR)^15,16^. It has been predicted that ~85% of firing rate variability, which can lead to decreased BCI performance, is due to physiological changes^8^. In the brain, the FBR is often characterized by a loss of neuronal density surrounding the implant, which has been shown in one report to decrease by 40% within 100μm^17^. Another prominent effect is the presence of reactive astrocytes surrounding the implant after 1-week post-implantation, shown by increased glial fibrillary acidic protein (GFAP)^15^. It has been shown that increased glial encapsulation can lead to increased impedance; however, it is not clear how this influences recording quality over time^9^. Furthermore, even though the FBR traditionally has been studied using the immunohistochemical (IHC) methods mentioned, neuron loss and GFAP intensity do not necessarily significantly predict signal stability^18^.

One promising area of research that could reveal biological mechanisms causing signal instability is the study of transcriptomics, which describes mRNA production (gene expression) of a biological system to gain insight on the biological processes being activated. This approach has been especially successful for studying diseases of the central nervous system (CNS) due to the incredible complexity and heterogeneity of cell types in the brain. For example, transcriptomics has revealed a novel target, neddylation, that ameliorates the severity of multiple sclerosis in a murine disease model^19^. Transcriptomics has also been applied in Alzheimer’s disease (AD) research to expose genetic, cell-type specific regulators of myelination that are perturbed in AD, as well as sex differences in the cellular response to AD^20^. Additionally, a transcriptomics method which allows for gene expression to be spatially resolved has been applied in AD research to reveal groups of genes that are differentially expressed surrounding amyloid plaques^21^.

Due to the power and novelty of transcriptomics to describe complex systems of aberrant physiological states, a new focus in the study of the brain-electrode interface is the transcriptomic profile of the FBR to implanted electrodes in the brain. Recent studies from our lab and others have illuminated specific genes that are differentially expressed around the implant compared to non-implanted brain tissue^22–24^. In our initial publication in this area, we used laser capture microscopy to dissect fixed tissue from near the device tract (<100μm), from tissue 500μm away from the device tract, and from a non-implanted animal. Differential expression analysis revealed 157 differentially expressed genes (DEGs) at 24 hours, 62 DEGs at 1 week, and 26 DEGs at 6 weeks post-implantation in comparison to non-implanted tissue. Differential expression (DE) analysis revealed several genes that were significantly differentially expressed between tissue 500μm from the device and naïve tissue, suggesting the spatial extent of differential gene expression extends past this distance. However, the spatial extent of differential gene expression remains unknown, as our previous analysis sampled discrete, pre-selected locations from the device interface.

Here, in an extension of our previous work^22^, we report the application of a newer spatial transcriptomics method which has three advantages over our previous approach: (1) it reports the transcriptional profile of individual genes with near cellular-scale resolution across the entire landscape of a tissue section containing the device, (2) it is likely to improve RNA quality compared to our previous work through the use of fresh-frozen tissue samples, and (3) it allows quantitative immunohistochemistry and spatial transcriptomics to be performed in the same tissue section. We applied this technique to the brain-electrode interface in rats implanted with single shank, silicon microelectrode arrays implanted in motor cortex. All timepoints (24-hour, 1-week, and 6-week implants) yielded significant, DEGs when comparing either implanted tissue sections to non-implanted tissue sections or an area ≤1 50μm from the device tract to an area ?500μm from the device tract within the same implanted tissue section. At 24 hours post-implantation, previously reported DEGs were expressed over a large portion of the tissue section, extending further than 3.0mm from the device tract. The expression of these genes consolidated around the device tract at 1 week and became even more compact at 6 weeks. Each timepoint revealed novel genes that have not been reported previously, and the spatial aspect of the data we present reveals important information about the FBR to implanted electrodes in the brain and its progression *in vivo.*

## II. Methods

### 2.1. Surgery, Implantation & Sample Preparation

All electrodes used for this study were single-shank, silicon, Michigan-style microelectrodes (A1×16-3mm-100-703-CMLP, 15 um thickness, NeuroNexus Inc, AnnArbor, MI). Adult, male, Sprague Dawley rats received implants in motor cortex using coordinates and procedures as previously described^22,25^. Briefly, electrodes were inserted (+3.0mm AP, 2.5mm ML from Bregma, 2.0 mm deep) during isoflurane anesthesia (~2.0% in oxygen), and the surgical site was closed via a dental acrylic headcap. Post-operative analgesia was achieved with injected meloxicam (2mg/kg) and topical application of bupivacaine. All animal procedures were approved by the Michigan State University Animal Care and Use Committee.

At the appropriate timepoint, the animals were euthanized via sodium pentobarbital administration and decapitation, and brains were removed and flash frozen with liquid nitrogen in optimal cutting temperature compound (OCT). Animals were not perfused prior to euthanasia to align with other reports^21,26,27^ and avoid any potential perturbation of gene expression during the process. Ten-micron thick cryosections were taken from primary motor cortex at a depth of ~1000μm and mounted on a Visium slide (10x Genomics, Pleasanton CA). Visium slides enable spatial transcriptomics through capture areas made up of an array of ~5,000 spots of spatially barcoded oligonucleotides which capture mRNA released from the tissue via a permeabilization step.

### 2.2. Staining & Imaging

Prior to tissue permeabilization, the tissue was methanol-fixed in cold methanol for 30 minutes, blocked in a bovine serum albumin (BSA) solution, and then labeled using a neuronal nuclei (NeuN) primary antibody (Rb pAB to NeuN, Abcam, Cambridge, MA, Cat #:104225) at a concentration of 1:100 and a GFAP primary antibody (Monoclonal Anti-glial fibrillary acidic protein (GFAP) antibody, Millipore Sigma, St. Louis, MO, Cat #: G3893-100) at a concentration of 1:400. Secondary antibodies used for staining were AlexaFluor 488 (Anti-rabbit IgG, Invitrogen, Eugene OR, Cat #: A11034) for conjugation to the NeuN primary and AlexaFluor 647 (Anti-mouse IgG, Invitrogen, Eugene, OR) for conjugation to the GFAP primary. Additionally, Hoechst (Life Technologies Corp. Eugene, OR) was used as a universal nuclei stain at a concentration of 1:1000. After staining, a Nikon A1R confocal microscope with a motorized stage was used to capture individual 20x magnification image tiles of each of the four capture areas on the Visium slide. The final wide-field image was created by software-automated stitching of the individual image tiles together.

### 2.3. Tissue Permeabilization & cDNA Synthesis

After imaging, the tissue is permeabilized using an enzyme for 18 minutes. This permeabilization time was established through an optimization experiment, where fluorescent complementary DNA (cDNA) was created on the slide to reveal the permeabilization time that maximized RNA release from a tissue section (**Fig. 1**). Although the fluorescence intensity appears similar between the 18- and 24-minute trials, the 24-minute trial showed evidence of over-permeabilization due to undefined cell body borders. Each capture area of a Visium Spatial Gene Expression Slide is filled with an array of ~5000, 55μm diameter spots each containing unique RNA-binding, spatially barcoded oligonucleotides (oligos) which capture mRNA released from the tissue for subsequent processing. After permeabilization and mRNA capture, reverse transcription is performed on the slide to extend the capture oligo based on the bound mRNA’s nucleotide sequence. The original mRNA strand is then released from the oligo and through a series of template switching and second strand synthesis steps, a final cDNA strand is produced containing both the mRNA sequence and spatial barcode. This cDNA strand is then released from the slide via denaturation and transferred to a DNA/RNA LoBind microcentrifuge tube (Eppendorf, Cat #: 022431021). The cDNA from each capture area is then quantified using qPCR and amplified using the number of cycles from qPCR required to achieve 25% of the peak fluorescence value. After amplification, the cDNA is purified using SPRIselect (Beckman Coulter Inc, Brea, CA), a paramagnetic bead-based size selection reagent.

**Figure 1:**
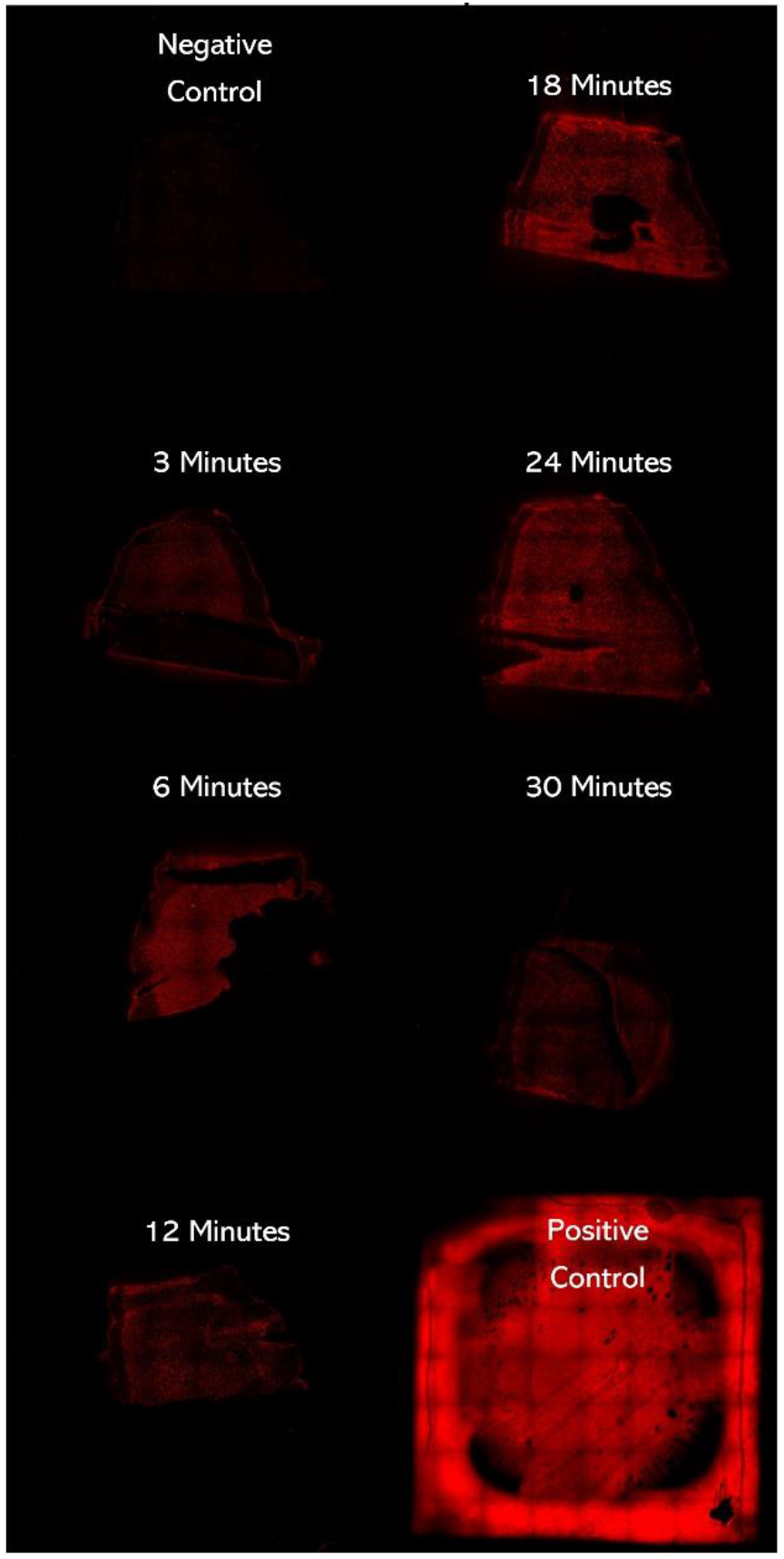
Permeabilization optimization experiments revealed 18 minutes as the appropriate incubation period for our studies. RNA was released onto a slide after the application of a tissue permeabilization enzyme after different timespans. The negative control (upper left) was not treated with permeabilization enzyme, and the positive control (lower right) received 2μL of reference rat RNA (RIN ≥ 7).

### 2.4 RNA Sequencing and Analysis

Samples were then transferred to the University of Michigan Advanced Genomics core for library preparation and sequencing. cDNA quality was assessed using the Tapestation 2200 (Agilent) and subjected to library preparation following the manufacturer’s protocol (10x Genomics). Final library quality was assessed using the LabChip GX (PerkinElmer). Pooled libraries were subjected to paired-end sequencing according to the manufacturer’s protocol (Illumina NovaSeq 6000). Bcl2fastq2 Conversion Software (Illumina) was used to generate de-multiplexed Fastq files and the SpaceRanger Pipeline (10x Genomics) was used to align reads and generate count matrices.

Sequencing results are in the form of raw .fastq files as well as a .cloupe file for each capture area that combines the spatial-barcode in the sequencing data to an image of the tissue section. This is made possible by a fluorescent fiducial frame that surrounds the capture area and is used as a reference to match the reads with unique spatial barcodes to their known location on the slide. The .cloupe file can then be loaded by 10x Genomics software, “Loupe Browser” which shows gene expression data overlaid on the image of each tissue section. Preliminary data shown in this report is produced from Loupe Browser analysis. Clusters of spots for differential expression analysis may be hand-selected and chosen to include any spots over the tissue. Log_2_(fold change), or LFC, is calculated between each cluster for each gene by the average number of counts of that gene detected from RNAseq. Counts of gene transcripts, or reads, within each spot are identified through RNAseq by a spatial barcode on each read while unique reads are identified through a unique molecular identifier (UMI). Therefore, a single count must have a unique UMI, spatial barcode, and gene annotation to be shown in either differential expression analysis or the spatial expression images below. DEGs are then used in gene ontology (GO) analysis, which attempts to take lists of DEGs and produce key terms that describe the active biological processes. The PANTHER Classification System^28^ is used for this purpose, and significant (p-value < 0.05) and highly enriched GO terms are reported for each timepoint. Specifically, the PANTHER Overrepresentation Test (Fisher’s exact test, Release: February 24, 2021) was used with a *Rattus norvegicus* reference from the GO Ontology database and GO biological process complete dataset for GO analysis (DOI: 10.5281/zendo.5228828, Release: August 8, 2021).

## III. Results and Discussion

### 3.1. Overview

This report details the use of a new spatial transcriptomics method that expands the current state-of-the-art in RNA-sequencing around electrode implants in multiple ways. First, the improved spatial sampling allows for a more complete understanding of the distance-based effects of the FBR to an implanted electrode. This is an advantage in our field because it allows spatial boundaries to be drawn around areas of interest for differential gene expression analysis. Moreover, the combination with immunohistochemistry within the same tissue sample reveals new information about the cellular source of individual genes driving the FBR. This will allow benchmarking newly identified biomarkers relative to traditional tissue response metrics within the same tissue section. Furthermore, avoiding the use of laser capture microscopy and paraformaldehyde fixation is expected to better preserve the RNA integrity number of these samples in comparison to our previous approach^22^.

While the data presented here should be interpreted with caution given the limited sample size (n = 1 rat per timepoint), they do point toward expression patterns that have not been shown previously and are a direct result of the additional capabilities of this spatial transcriptomic method. When compared to a non-implanted naïve tissue section, differential expression analysis revealed 5811 genes in the 24-hour sample, 2422 genes in the 1-week sample, and 513 genes in the 6-week sample (p-value < 0.05). When differential expression analysis was performed between an area near the device tract (≤1 50μm) and an area far from the device tract (≥500μm), it yields 1056 genes in the 1-week sample, and 163 genes in the 6-week sample (p-value <0.05)(this comparison was omitted for the 24 hour sample, as described below). When the comparison to naïve tissue is used, there is a general decrease in the number of differentially expressed genes over time, post-implantation. This trend has been described in our previous work as well which found 157, 62, and 26 DEGs compared to a naïve tissue section at 24 hours, 1 week, and 6 weeks post-implantation, respectively^22^. Other work has also shown that differential gene expression is highest at 24 hours compared to 6 hours, 3 days, and 2 weeks^23^. Each timepoint and tissue section from these new experiments will now be explored more in detail to show unique timepoint-based effects on both the spatial and transcriptional identity of differential gene expression surrounding an implanted electrode.

### 3.2. 24 Hours

An immediate observation, which is made possible by the spatial aspect of this method, is that the spatial expression pattern of DEGs at 24 hours, compared to a naïve tissue section, extends far past 500μm (**Fig. 2**). For many genes, expression extends up to the edge of the tissue section more than 3.0mm away from the device tract (**Supplemental Fig. 1**). *Gfap*, a prototypic marker of astroglial reactivity to implanted devices, is one of the more prominent examples of this large spatial footprint (**Fig. 2C**). However, compared to non-implanted tissue, vimentin (*Vim*) and glycoprotein nonmetastatic melanoma protein B (*Gpnmb*) are also differentially and broadly expressed (**Fig. 2D-E**). The spatial breadth of the observed changes in gene expression at 24 hours led us to forego a within-tissue, distance-based comparison for differential expression in favor of a whole tissue section comparison to non-implanted, naïve tissue. Using this comparison for differential expression analysis, the top 75 DE genes with the largest LFC are shown in the **Fig. 2 Table**. The top three enriched GO terms for the 5811 identified DE genes using the PANTHER Classification System are “regulation of long-term neuronal synaptic plasticity” (fold enrichment: 3.06, p-value: 1.28E-5), “myelin assembly” (fold enrichment: 2.87, p-value: 1.13E-3), and “synaptic vesicle priming” (fold enrichment: 2.83, p-value: 3.08E-3). The top three most significant differentially expressed genes that relate to “regulation of long-term neuronal synaptic plasticity” are regulating synaptic membrane exocytosis protein 1 (*Rims1*, LFC: −2.895, p-value: 5.84E-76), Alpha-synuclein (*Snca,* LFC: −1.378, p-value: 1.40E-31), and Ras/Rap GTPase-activating protein SynGAP (*Syngap1*, LFC: −2.687, p-value: 8.22E-16). The top three DEGs that relate to “myelin assembly” are myelin-associated oligodendrocyte basic protein (*Mobp*, LFC: −1.896, p-value: 1.41E-40), glypican-1 (*Gpc1*, LFC: −1.126, p-value: 1.04E-15), and Ankyrin 2 (*Ank2*, LFC: −1.033, p-value: 4.33E-13). The top three DEGs genes found that relate to “synaptic vesicle priming” are *Rims1, Snca,* and synaptojanin-1 (*Synj1,* LFC: −1.02, p-value: 2.57E-17). Negative LFCs of the genes related to “regulation of long-term neuronal synaptic plasticity” and “myelin assembly” indicate a breakdown of these normal homeostatic mechanisms in the tissue at 24 hours post-electrode implantation. Recent works have implicated both of these mechanisms in disruption of the normal function of tissue surrounding electrode implants^22,29^. Reduced expression of genes related to “synaptic vesicle priming” may be related to a response to excitotoxicity at this early timepoint.

**Figure 2:**
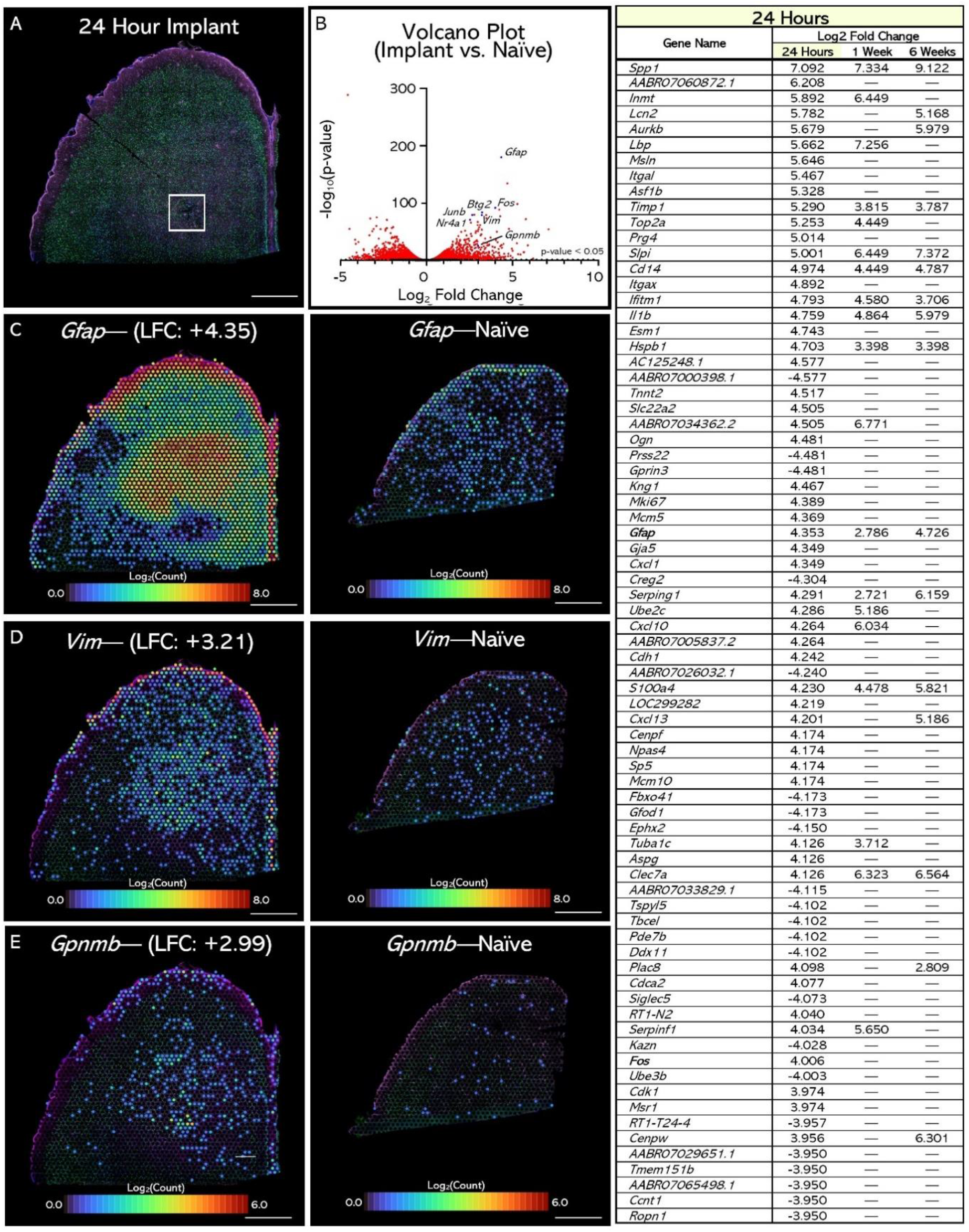
Spatial transcriptomics reveals changes in gene expression in primary motor cortex of a rat implanted for 24 hours. **(A)** IHC image of the tissue section (green: NeuN, magenta: GFAP, blue: Hoechst). White box shows implant site. **(B)** Volcano plot of differentially expressed genes between the 24-hour implant tissue section and a non-implanted, naïve tissue section taken from rat motor cortex (not shown). **(C, D, E)** Spatial expression pattern of genes previously reported in transcriptomic studies of the FBR to implanted electrodes. Left column shows gene expression in the 24-hour sample and right column shows gene expression in a naïve sample. **(Table)** Top 75 DE genes in the 24-hour comparison and those genes’ DE in the 1- and 6-week sample comparisons. Bolded genes denote genes highlighted in panels C-E and Fig. 3. Scale bars are 1000μm.

DE analysis revealed genes that have implications on tissue function surrounding the implanted electrode. **Fig. 3** highlights four of these, which were selected based on their possible role in FBR initiation and synaptic function. The first and second, *Fos* and *Junb,* are members of a family of “inducible transcription factors” that dimerize to form an activator protein 1 (AP-1) complex which serves as a transcription factor for genes that become activated in response to diverse forms of stimuli^30^. Differential expression of *Fos* and *Jun* genes and their AP-1 complex has been extensively described in many brain injury models, from traumatic brain injury^31^ to irradiation of brain tissue^32^ and electroconvulsive seizures^33^. AP-1 has been found in brain injuries leading to excitotoxicity and programed cell death^30^. Previous work from our lab^25^ as well as this study (**Supplemental Fig. 2**) show changes in ion channel expression, possibly leading to hyperexcitability and excitotoxicity at early timepoints. These findings, combined with *Fos/JunB/*AP-1 activation soon after electrode implantation, could align with an excitotoxic mechanism leading to neuronal cell death following insertional damage.

**Figure 3:**
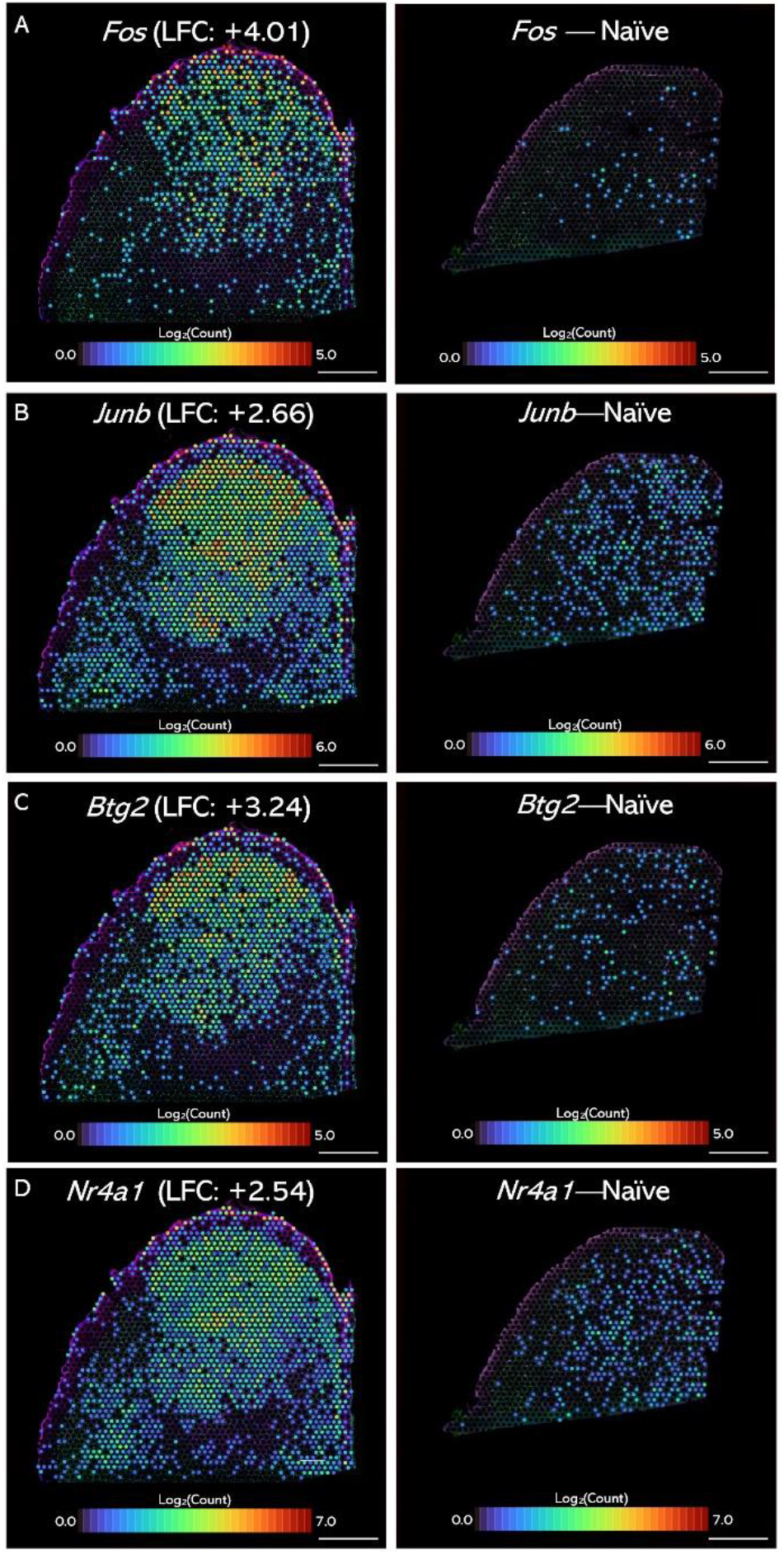
Spatial expression patterns of differentially expressed genes in the 24-hour (left) and naïve (right) samples. LFC is calculated between the 24-hour and naïve whole tissue sections. Scale bar: 1000μm

The third gene highlighted in **Fig. 3C** is *Btg2,* which encodes the protein “B-cell Translocation Gene 2.” Btg2 is considered an anti-proliferative protein which arrests the cell cycle and may prevent apoptosis in neurons^34,35^. However, one study finds that inhibiting the action of Btg2 via microRNA-146 reduces apoptosis^36^, and another shows that minocycline, a drug with neuroprotective effects, increases the expression of Btg2^37^. In glia, deletion of *Btg2* increases the amount of GFAP, a common reactive astrocyte marker, in a bilateral common carotid artery stenosis model, emphasizing the anti-proliferative role of Btg2^38^. While the exact effect of increased Btg2 expression *in vivo* after brain injury is debatable, the finding of increased expression of *Btg2* in the FBR to implanted electrodes aligns our results with these other injury models.

The fourth and final gene highlighted in **Fig. 3D** is *Nr4a1*, which encodes the nuclear transcription factor “Nuclear Receptor Subfamily 4 Group A Member 1.” Continuing with the theme of a transcriptional response to excitotoxicity, *Nr4a1* is dramatically increased following induced seizures^39^. However, it is also commonly reported in studies of neuroinflammation^40,41^ and stress^42^. In Estrada et al., *Nr4a1* knockout mice treated with lipopolysaccharide (an inducer of neuroinflammation) showed reduced amount of inflammation compared to wild-type mice. Specifically, the *Nr4a1* knockout animals had a decreased amount of activated microglia (measured by Iba1+ cells), and inflammatory cytokines *Il1b*, *Il6* and *Tnf*. Another study by Jeannetau et al. shows a similar protective effect of *Nr4a1* knockout in a murine chronic stress model. They also found that *Nr4a1* over-expression decreased the number of spines on the apical tuft dendrites of pyramidal neurons *in vivo* and that this decrease required *Nr4a1* expression. Loss of dendrites and dendritic spines around electrode implants is a current area of study^29^ and is another example of the FBR causing changes in the structure and function of the surrounding neural parenchyma. These effects of *Nr4a1* on dendrites contribute to the regulation of synaptic plasticity, which shows up as a GO term for the 24-hour timepoint. Neuronal *Nr4a1* also upregulates genes related to mitochondrial uncoupling, thereby decreasing the amount of reactive oxygen species generation by mitochondrial ATP production^43^. It is possible that *Nr4a1* upregulation is a compensatory mechanism by the brain in response to the damaging effects of an electrode implant.

### 3.3. 1 Week

At 1-week post-implantation, *Gfap*, *Vim*, and *Gpnmb* expression consolidated around the device tract and resembled a pattern similar to that reported in literature, allowing for a ≤150μm vs ≥500μm comparison to measure gene expression differences most relevant to the electrophysiological function of the implanted device. One unexpected observation that becomes more prominent at 1 week is that the glia limitans (arrow in **Fig. 4A**) differentially expresses many of the genes found to be differentially expressed within 1 50μm of the device tract. This observation will be discussed in more detail later; however, due to this observation, the glia limitans was removed from differential expression analysis in the comparison of tissue ≤150μm to tissue ≥500μm. In total, this analysis revealed 1056 DEGs at the brain-electrode interface. In the **Fig. 4 Table** the top 75 DE genes with the largest LFCs are shown. In gene ontology, the top three enriched GO terms using the PANTHER Classification System are “immune complex clearance” (fold enrichment: 20.25, p-value: 1.87E-3), “positive regulation of inflammatory response to wounding” (fold enrichment: 20.25, p-value: 1.87E-3), and “complement-mediated synapse pruning” (fold enrichment: 20.25, p-value: 1.87E-3). The genes found that relate to immune complex clearance are complement factor H (*Cfh*, LFC: +2.50, p-value: 4.51E-12), Clusterin (*Clu*, LFC: +1.28, p-value: 2.83E-4), and low affinity immunoglobulin gama Fc region receptor II (*Fcgr2b*, LFC: +3.00, p-value: 2.17E-6). The genes found that relate to the positive regulation of inflammatory response to wounding are signal transducer CD24 (*Cd24*, LFC: +1.21, p-value: 0.0339), progranulin (*Grn*, LFC: +2.83, p-value: 3.99E-18), and midkine (*Mdk*, LFC: +1.73, p-value: 2.34E-4). The genes found that relate to complement-mediated synapse pruning are compliment C1q subcomponent subunit A (*C1qa*, LFC: +3.14, p-value: 8.37E-25), IG domain-containing protein (*Trem2*, LFC: +3.08, p-value: 1.25E-14), and complement C3 (*C3*, LFC: +1.17, p-value: 0.030). Immune and inflammation-related pathways are expected at 1-week post-implantation^15,22,24^.The final GO term, “compliment-mediated synapse pruning” is especially interesting as it relates to recent findings from our lab relating to changes in ion channel expression and hyperexcitability of neural tissue surrounding electrodes immediately following implantation^25^ as well as even more recent findings that structural remodeling of dendritic spines occurs following implantation^29^.

**Figure 4:**
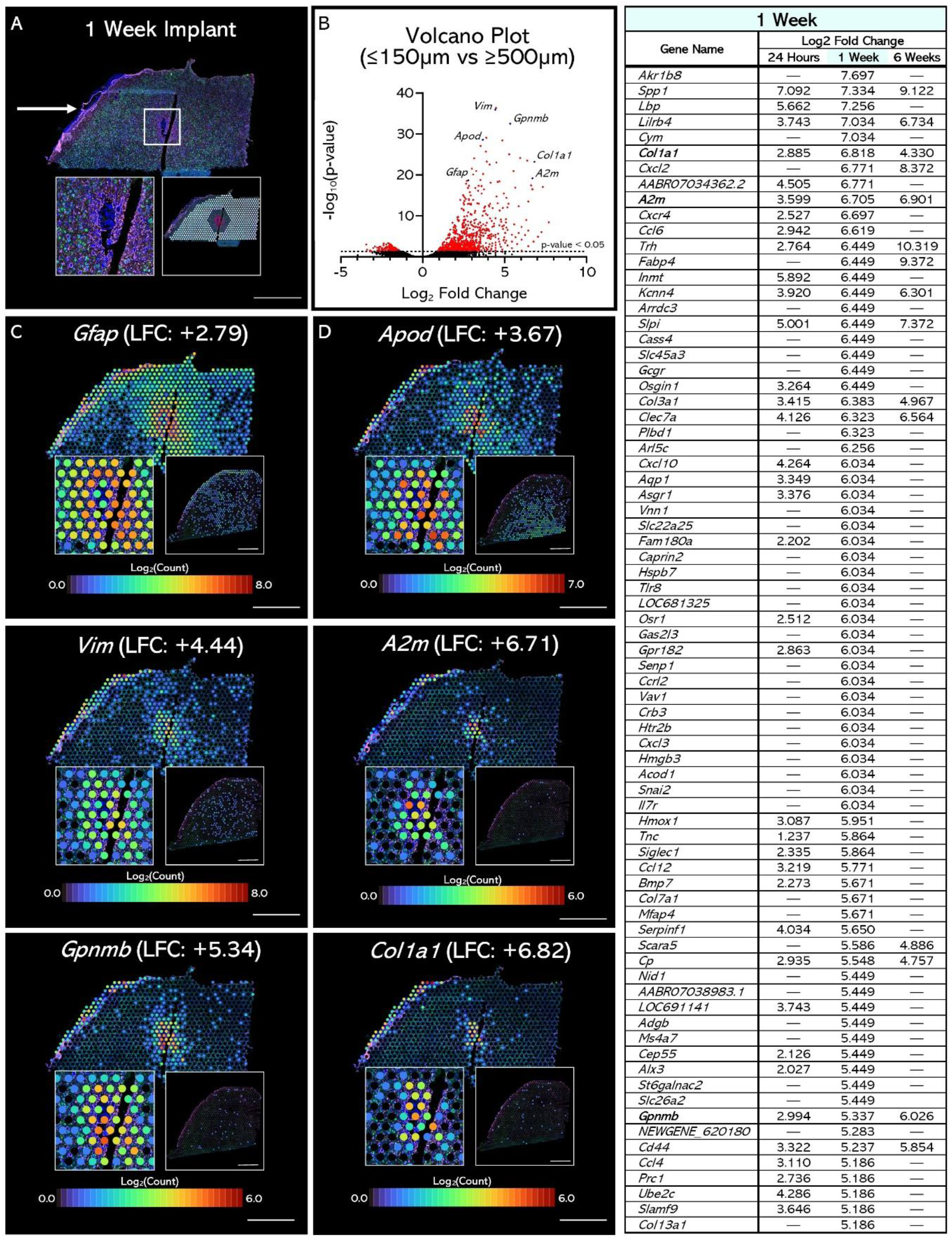
Spatial transcriptomics reveals changes in gene expression in primary motor cortex of a rat implanted for 1 week. **(A)** IHC image of the tissue section (green: NeuN, magenta: GFAP, blue: Hoechst). White box shows implant site. **(B)** Volcano plot of differentially expressed genes between an area within 150μm of the electrode tract and an area greater than 5øøμm from the electrode tract, excluding the glia limitans. **(C, D)** Spatial expression pattern of previously reported genes (C) and novel genes (D). Left inlay shows zoomed in image of the electrode tract. Right inlay shows the gene’s expression in a naive tissue section. **(Table)** Top 75 DE genes in the 1-week comparison and those genes’ DE in the 24-hour and 6-week sample comparisons. Bolded genes denote genes highlighted in panels C-D. Scale bars are løøøμm.

DE analysis of the 1-week tissue revealed 1056 genes. Due to the GO terms for these genes generally relating to an activated immune response and inflammation, **Fig. 4D** highlights three genes that have been found to play a role in these mechanisms. The first, Apolipoprotein D (*Apod*) is upregulated in the brain in Alzheimer’s disease, especially in the late disease stages and in oligodendrocyte precursor cells^20,21,44,45^. The general function of ApoD protein is lipid transport and it has been shown to exert neuroprotective effects by reducing the harmful effects of oxidative stress^44,46^. In the context of a FBR, this could indicate the activity of a homeostatic mechanism by which the surrounding tissue is repaired and the damaging effects of the inflammatory response are mitigated.

The next gene shown in **Fig. 4D** is Alpha-2-macroglobulin (*A2m*), a major component of the innate immune system that mediates transferrin binding, serves as a protease inhibitor, and may contain inflammation in the brain by scavenging myelin basic protein^47,48^. Both transferrin and myelin basic protein are upregulated around the device at 1 week in this study, and these genes have been shown in our previous transcriptomics study as well^22^. A2M is highly upregulated in Alzheimer’s disease, colocalizing with amyloid plaques^49^, as well as in multiple sclerosis, brain trauma, bacterial meningitis, and cerebral infarction^50^. The activity of A2M in biological function is widespread; however, it largely seems to serve a protective role against inflammation. A2m is also upregulated in the 24-hour sample (LFC: +3.599, p-value: 1.93E-27) and the 6-week sample (LFC: +6.90, p-value: 1.59E-7).

The final gene highlighted in **Fig. 4D** is Collagen1A1 (*Col1a1*), which encodes a subunit of collagen protein. It is commonly found in connective tissue; however, it has been found in perivascular stromal cells (PSCs) that form the fibrotic scar in CNS injury^51^. After ischemic stroke for example, there is a proliferation of these perivascular stromal cells that forms a fibrotic scar at the core of the ischemic lesion within the boundaries of the astroglial response^52^. This finding emphasizes that the “glial scar” surrounding device implants is only part of a larger process surrounding the device with fibrous tissue, and that these other cell types may be influencing the surrounding tissue in different ways. For example, it has been shown that Col1a1 positive PSCs produce retinoic acid, a signaling molecule that regulates gene expression throughout the body^51^.

### 3.4. 6 Weeks

At 6 weeks post-implantation, *Gfap, Vim,* and *Gpnmb* are all differentially expressed within 1 50μm of the device tract compared to the area greater than 500μm from the device tract, excluding the glia limitans. The trend of consolidating spatial extent of the DEGs continues at 6-weeks, shown by a much smaller radius of most of the genes found compared to 24 hours or 1 week. There are also far fewer differentially expressed genes at 6 weeks compared to 24 hours and 1 week, a finding consistent with other reports^24^, indicating the FBR reaches a kind of homeostasis in the chronic phase. Still, the brain-electrode interface is highly active at 6 weeks, differentially expressing 163 genes. In the **Fig. 5 Table** the top 75 DE genes are shown with the largest LFCs.

**Figure 5:**
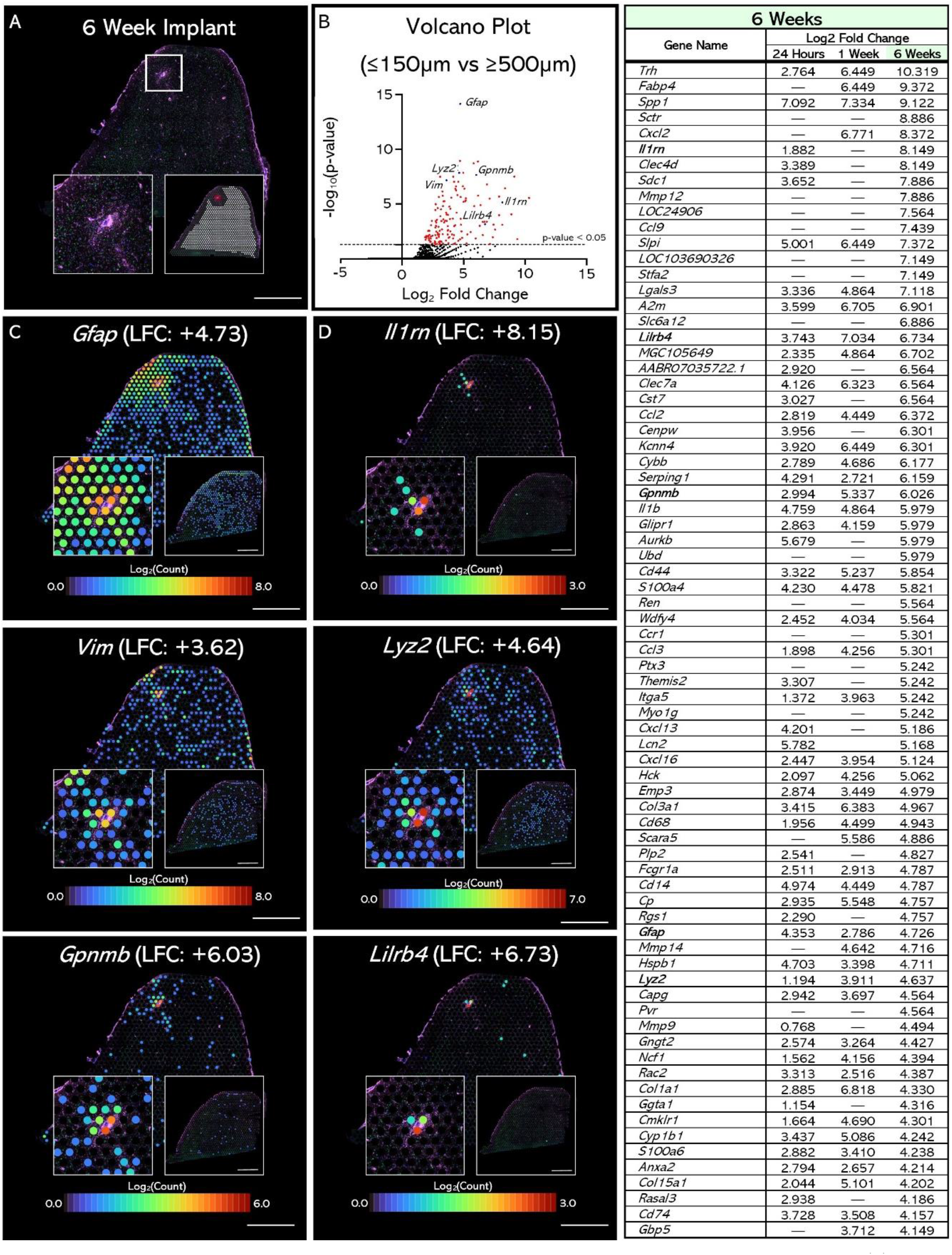
Spatial transcriptomics reveals changes in gene expression in primary motor cortex of a rat implanted for 6 weeks. **(A)** IHC image of the tissue section (green: NeuN, magenta: GFAP, blue: Hoechst). White box shows implant site. **(B)** Volcano plot of differentially expressed genes between an area within 150μm of the electrode tract and an area greater than 500μm from the electrode tract, excluding the glia limitans. **(C, D)** Spatial expression pattern of previously reported genes (C) and novel genes (D). Left inlay shows zoomed in image of the electrode tract. Right inlay shows the gene’s expression in a naive tissue section. **(Table)** Top 75 DE genes in the 6-week comparison and those genes’ DE in the 24-hour and 1-week sample comparisons. Bolded genes denote genes highlighted in panels C-D. Scale bars are 1000μm.

At 6 weeks, the top three enriched gene ontology (GO) terms from the differentially expressed genes are “negative regulation of complement activation, lectin pathway” (fold enrichment: >100, p-value: 3.32E-4), “Fc-gamma receptor signaling pathway involved in phagocytosis” (fold enrichment: >100, p-value: 3.32E-4), and “vertebrate eye-specific patterning” (fold enrichment: >100, p-value: 3.32E-4). The genes found that relate to the “negative regulation of complement activation, lectin pathway” are plasma protease C1 inhibitor (*Serping1*, LFC: +6.16, p-value:1.30E-9) and alpha-2-macroglobulin (*A2m*, LFC: +6.90, p-value: 1.59E-7). The genes found that relate to “Fc-gama receptor signaling pathway involved in phagocytosis” are myosin IG (*Myo1g*, LFC: +5.24, p-value: 0.0391) and low affinity immunoglobulin gamma FC region receptor II (*Fcgr2b*, LFC: +3.61, p-value: 8.83E-5).

The first GO term, relating to a “negative regulation of complement activation,” could possibly indicate an attempt to maintain homeostasis at the interface by an active repression of the FBR. For this reason, the three genes we chose to highlight in **Fig. 5D** have been reported to negatively regulate either inflammation or the immune response. The first gene highlighted in **Fig. 5D** is *Il1rn* which encodes the protein “interleukin 1 receptor antagonist”. Il1rn is acts as an inhibitor of interleukin 1 (IL-1), a cytokine with a central role in the inflammatory response to brain injury, by competing with IL-1 for the binding site on the IL-1 receptor^53^. Systemic delivery of Il1rn after traumatic brain injury has been shown to reduce neuronal cell death and improve cognitive function after traumatic brain injury, indicating Il1rn has a neuroprotective effect when delivered in a neuroinflammatory environment^54^. Il1rn has also been shown to reduce systemic inflammation markers in humans after subarachnoid hemorrhage.^55^ The presence of *Il1rn* expression at 6 weeks post-implantation (but not at 24 hours or 1 week) indicates a movement towards an anti-inflammatory transcriptomic profile as time progresses, possibly to regain homeostasis by inhibiting the inflammatory cascade. While it is not one of the listed genes related to the “negative regulation of complement activation, lectin pathway” GO term, *Il1rn* does fit into the paradigm of a movement towards a negative regulation of the overall inflammatory response.

The second gene highlighted in **Fig. 5D** is *Lyz2* which encodes lysozyme, a component of the innate immune response that is usually thought to be found in macrophages, microglia, and astrocytes in the brain but has recently been found in retinal neurons as well^56,57^. Lysozyme is elevated in the cerebrospinal fluid of patients suffering from CNS infection^58^ and is generally implicated in an anti-inflammatory role by acting against oxidants^59^, reducing the expression of inflammatory cytokines^60^, and inhibiting the complement cascade^61^. Lysozyme is also elevated in the cerebrospinal fluid of patients suffering from Alzheimer’s disease and is colocalized with amyloid plaques. In *in vitro* studies, lysozyme has been shown to prevent aggregation of amyloid plaques^62^. It is possible that lysozyme is acting in a similar fashion in the case of the FBR to implanted electrodes by breaking down the fibrous scar formed around the implant as well as acting as a general inhibitor of inflammation, further emphasizing the anti-inflammatory profile of the interfacial tissue 6 weeks post-implantation.

The final highlighted gene in **Fig. 5D** shows the spatial expression pattern of *Lilrb4*, a gene that encodes an immune inhibitory immunoglobulin-like protein, Leukocyte immunoglobulin-like receptor B4. This receptor serves an important and complex role in the function of the immune system. LILRB4 is expressed in many different types of immune cells such as T-cells, granulocytes, dendritic cells, macrophages, monocytes, B cells, natural killer cells, mast cells, and microglia. LILRB4 acts as a regulator of the immune response in healthy systems to prevent autoimmune disorders; however, it also participates in many disease states and is over-expressed in both microglia around amyloid plaques in Alzheimer’s disease^63^ and in peripheral blood monocytes in multiple sclerosis patients^64^. In these neurological disorders, LILRB4 is generally thought to play an immune-suppressive role to mediate the immune response. Therefore, its presence at the brain-electrode interface indicates that, along with the anti-inflammatory aspects previously described, there is also active suppression of the immune response 6 weeks post-implantation.

### 3.5. Anomalies

#### I. Spatial Patterns of Gene Expression at 24 Hours

When the spatial pattern of previously reported DEGs such as *Gfap*, *Vim*, and *Gpnmb* is explored in the 24-hour sample, the spatial footprint is not centered around the device tract (**Supplemental Fig. 1C**) but appears to be centered around an area ~1.5mm anterior from the device tract. Additionally, the genes that are differentially expressed between the 24-hour implant sample and a naïve sample also surround this area and not the device tract. Some examples are ion channel/pump related genes and immediate early genes (**Supplemental Fig. 1**). After closer inspection, the IHC image of the tissue in this area shows a staining artifact consistent with bleeding (**Supplemental Fig. 1B**). There is another similar staining artifact found ~1.8mm lateral of the device tract that differentially expresses some genes, but not to the same extent as the first. This could indicate that the immediate inflammatory/immune response occurs from blood vessel rupture and that over time (after 24 hours, but before 1 week) the inflammatory response moves to surround the electrode implant.

Another unexpected observation of gene expression at 24 hours is that, in general, expression is greatly reduced immediately surrounding the device tract. There are many reports documenting cell death surrounding electrodes post-implantation which could be the cause of this decrease in transcription. However, the size of reduced transcription for many genes (ex. >500μm diameter for *Gfap,* **Fig. 2C**) is much larger than the area usually reported for cell death^65^. Another possible explanation for this observation would be that damage caused by implantation somehow interferes with transcription, and the inflammation/immune response that follows succeeds in repairing the tissue by 1 week to somewhat normal function. Another oddity of this observation is that IHC staining of tissue surrounding the device tract shows intact nuclei with no remarkable disruptions that one would expect from extensive damage causing transcriptional disruption. One final explanation is that there is a limitation with this method due to a difference in tissue permeabilization caused by electrode implantation. Due to the unremarkable nuclei staining in this section and even permeabilization in the optimization experiment, this explanation is less likely, and the relatively reduced gene expression surrounding the electrode implant at 24 hours may be a real, biological effect. Additional samples will be assessed to determine whether or not this effect is repeatable across animals.

#### II. Glia Limitans

The glia limitans^66^ forms a barrier between the meninges and neural parenchyma. It is similar to the blood-brain-barrier, as well as to glial scarring, which is an important part of the FBR to implanted electrodes. These barriers serve to protect the neural parenchyma by forming an “immune privileged” area in the brain, restricting leukocyte and other inflammatory cells from migrating across the barrier. The glia limitans, located on the outer edge of each tissue section (arrow in **Fig. 4A**), differentially expresses many of the same genes found near the device tract, especially in the 1-week tissue section. For this reason, the glia limitans was removed from differential expression analysis because its spatial position greater than 500μm from the device contributes to the cluster of spots used as a control for the 1 50μm vs. 500μm DE comparison.

To the best of our knowledge, broad inflammation in the glia limitans following implantation is a novel observation for cortical implants, and has inspired a new hypothesis: the transcriptomic effects of implantation may not only extend down the electrode tract, but also across the cortical surface. What are the implications of this observation? The literature available to interpret this effect is relatively limited, with a notable exception of a recently published overview of astrocytic barriers^65^. It is possible that the glia limitans and device-associated glial encapsulation might be functioning as a single tissue, essentially amplifying the inflammatory response. While it is not yet clear how differential expression of inflammatory genes in the glia limitans affects the surrounding tissue, if glia limitans inflammation does affect the structure or function of the adjacent neural parenchyma, the effects of the FBR to implanted electrodes could extend distances across the surface of the cortex orders of magnitude greater than previously reported. This could be an important consideration in the case of multiple implants, particularly if the insertions are performed during separate surgeries.

Regardless of the mechanisms causing glia limitans inflammation or their effects on the function and structure of surrounding tissue, future studies of the FBR to implanted electrodes should consider effects driven not just by the local response surrounding the electrode implant, but also a more global response from the glia limitans. To understand the role of the glia limitans in the FBR, future studies will need to address whether glia limitans inflammation exerts any effects on the structure or function of the surrounding neural tissue, since inflammation in the glia limitans extends far past the inflammatory effects at the electrode interface. Also, from these experiments we cannot fully assess the spatial extent of differential gene expression in the glia limitans; therefore, it will be important in future studies to explore exactly how far the inflammatory DEGs extend in the glia limitans.

#### III. Conclusion

Using spatial transcriptomics to study the FBR to implanted electrodes for the first time, these results reveal new information by measuring gene expression as a function of location and distance relative to the electrode tract, as well as using a precise collection method to measure gene expression in specific areas relevant to the function of recording electrodes. Comparing gene expression in the 150μm radius surrounding an implanted electrode to the area of tissue greater than 500μm from the tract (excluding the glia limitans) yields 1056 DE genes in the 1-week sample, and 163 DE genes in the 6-week sample. This yield is many times greater than the number of DE genes found from similar comparisons in our previous transcriptomics work. With the addition of spatial information, new observations were made possible, such as broad expression of inflammatory markers in the glia limitans and the complex spatial pattern of inflammatory markers 24 hours post-electrode implantation. Regarding the 24-hour section, the comparison to naïve tissue yielded 5811 DE genes, some of which relate to cellular mechanisms of excitotoxicity and an early, acute response to stimuli. At 1-week post-implantation, many classical markers of the inflammatory response and complement cascade were found. Finally at 6 weeks, genes that relate to the inhibition of inflammation are found, indicating a move towards homeostasis and depression of the FBR in the chronic phase.

Future work on this project will include additional sample collection to supplement the data outlined in this paper. Further data collection will also be complemented by electrophysiological data analysis. The goal of analyzing electrophysiological as well as gene expression and IHC data will be to find relationships between traditional biomarkers of the FBR, recording quality and stability, and gene expression. In order to accomplish this, gene expression data must be simplified, and the most important effects drawn out. To this end, we will apply computational analysis techniques to the spatial transcriptomics data in future studies^21^. Transcriptomics has been successfully applied in other fields to gain new insight into longstanding biological mysteries. We hope that with the additional capabilities of spatial transcriptomics, this new method will be a step towards a deeper understanding of the brain-electrode interface and the FBR to implanted electrodes in the brain.

## Supporting information

Differential Gene Expression Analysis

## Acknowledgments

The authors thank Dr. Oliva Koues and Grace Kenny at the University of Michigan Advanced Genomics core for RNA sequencing and initial data processing. Dr. Melinda Frame, academic specialist at the Center for Advanced Microscopy at Michigan State University, provided valuable support setting up confocal imaging for these experiments. Cort Thompson provided helpful feedback and discussions throughout experimentation and manuscript preparation. Dr. Evon S. Ereifej also provided valuable feedback on the content of this manuscript.

---

**Supplemental Figure 1:**
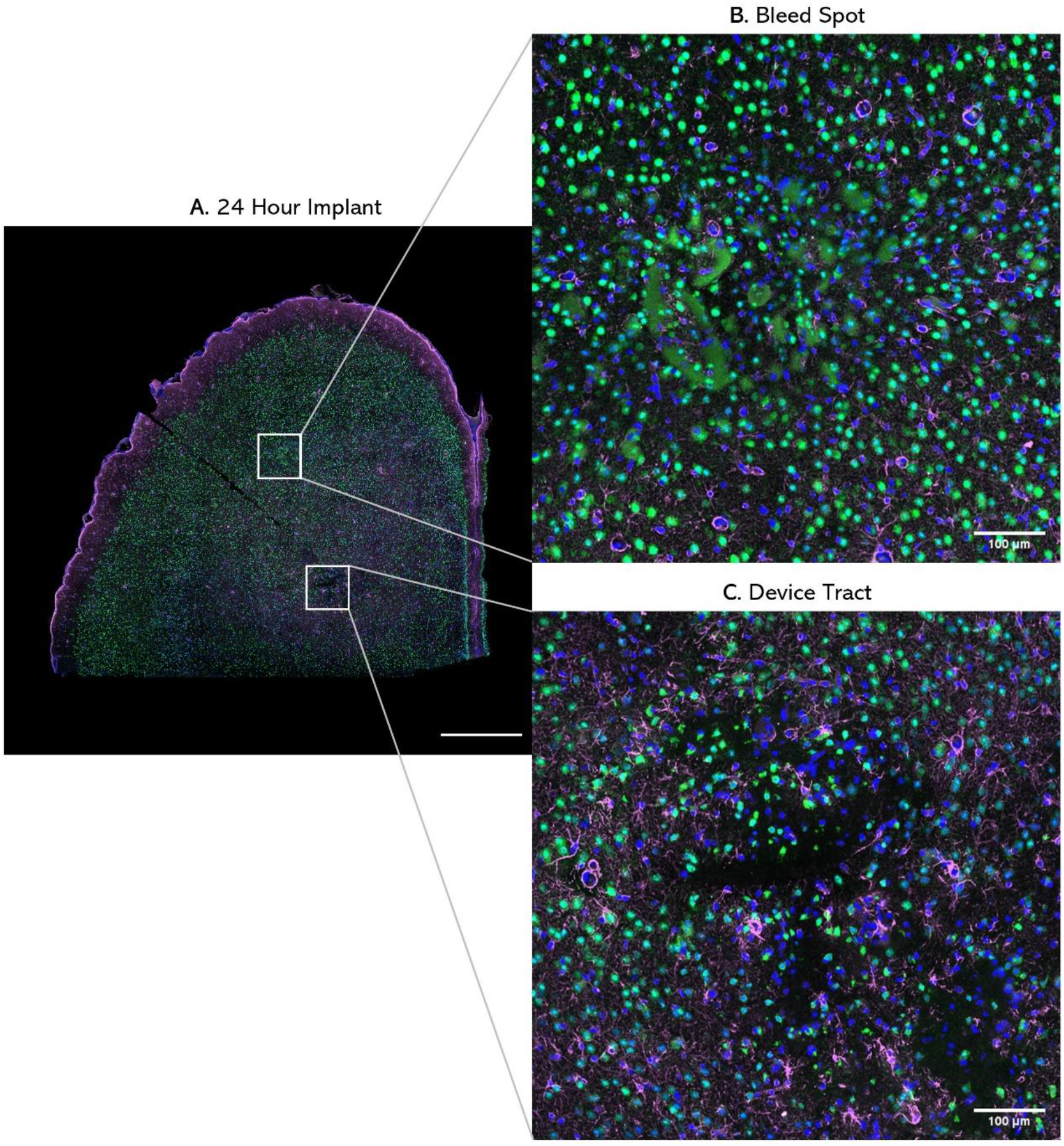
**(A)** IHC image of a 10μm thick tissue section implanted with an electrode for 24 hours. Scale bar is 1000μm. **(B)** Zoomed in image of an IHC staining artifact which differentially expresses many genes related to the FBR. **(C)** Zoomed in image of the electrode tract. Unlabeled scale bar: 1000μm

**Supplemental Figure 2:**
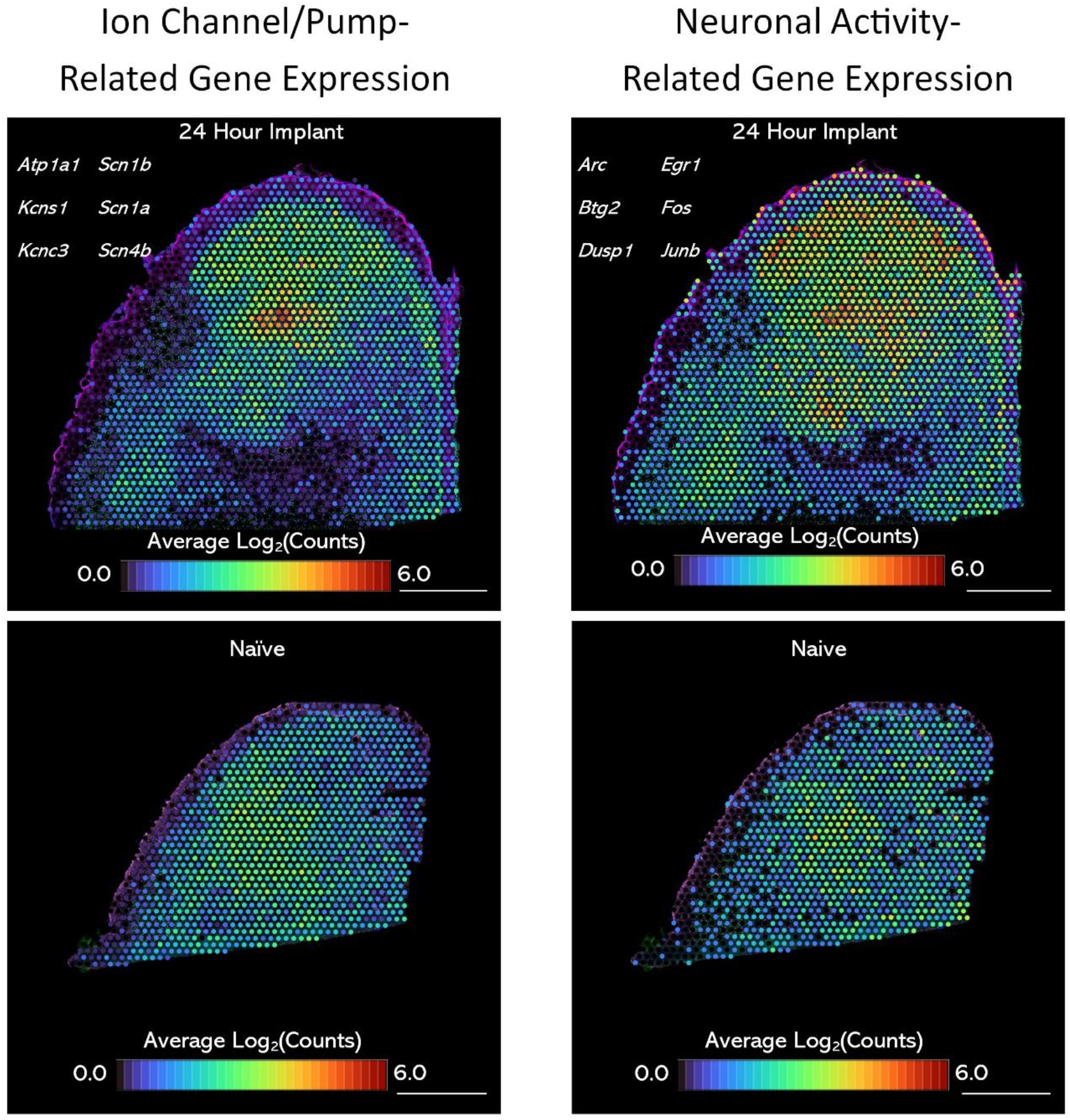
Spatial expression patterns of genes related to ion channels/pumps (left) and immediate early genes (right) in a tissue section implanted with an electrode for 24 hours (top) and a naïve tissue section (bottom). Scale bars are 1000μm.

